# Comparative Characterization Reveals Conserved and Divergent Ecological Traits of Oral Corynebacteria

**DOI:** 10.1101/2025.09.16.676652

**Authors:** Molly Burnside, Emily Helliwell, Puthayalai Treerat, Tanner Rozendal, Justin Merritt, Jonathon L. Baker, Jens Kreth

## Abstract

Corynebacteria are abundant members of the oral microbiome and increasingly recognized as key structural organizers of supragingival biofilms. Despite their prevalence, the ecological roles and phenotypic traits of many oral corynebacterial species remain poorly defined. Here, we isolated and characterized two new strains, *Corynebacterium durum* JJ2 and *Corynebacterium argentoratense* MB1, and compared them with reference strains *Corynebacterium durum* JJ1 and *Corynebacterium matruchotii* ATCC14266. Phenotypic assays revealed that *C. durum* strains displayed robust aggregation, thick biofilm formation, and extensive extracellular polymeric substance (EPS) networks, whereas *C. argentoratense* and *C. matruchotii* formed thinner biofilms with minimal EPS production. All four strains secreted extracellular membrane vesicles (EMVs) capable of inducing chain elongation in *Streptococcus sanguinis*, underscoring a conserved interspecies signaling function. Genomic analysis demonstrated close relatedness between *C. durum* and *C. matruchotii*, while *C. argentoratense* was more distantly related, with a reduced genome, fewer metabolic pathways, and absence of nitrate reductase genes, consistent with its inability to grow under anaerobic conditions. These findings suggest that *C. argentoratense* may represent a less specialized or transient inhabitant of the oral cavity, whereas *C. durum* and *C. matruchotii* are well adapted to the oral niche. Together, this study expands our understanding of phenotypic diversity, metabolic capacity, and interspecies interactions among oral corynebacteria, highlighting their potential importance as biofilm organizers and contributors to oral microbial ecology.

**Importance:** Oral Corynebacteria contribute to the structural and ecological stability of supragingival communities. Yet, their species-level functions remain poorly defined. By isolating and characterizing new strains of *Corynebacterium durum* and *Corynebacterium argentoratense*, and comparing them with reference strains including *C. matruchotii*, we provide new insight into their phenotypic diversity, metabolic capacity, and ecological roles. Our results demonstrate that *C. durum* strains form robust biofilms enriched in extracellular polymeric substances, while *C. argentoratense* produces thinner biofilms and lacks the genomic features required for anaerobic growth, suggesting a less specialized or transient role in the oral cavity. Importantly, we show that extracellular membrane vesicles secreted by all tested strains promote chain elongation in *Streptococcus sanguinis*, highlighting a conserved mechanism of interspecies communication. These findings advance our understanding of how oral corynebacteria contribute to biofilm organization and microbial homeostasis, and position them as critical but understudied players in oral microbial ecology.

## Introduction

The human oral microbiome is one of the most densely populated and taxonomically diverse microbial communities in the body (1). The expanded Human Oral Microbiome Database (eHOMD) currently lists over 800 microbial taxa associated with the oral cavity (2). While each individual harbors a subset of approximately 150 to 300 species, the composition of this community plays a pivotal role in determining the status of oral health and disease (3). Importantly, many oral disease-associated microorganisms do not meet the classical criteria of pathogens as defined by Koch’s postulates. Instead, these organisms referred to as pathobionts can transition from harmless to harmful depending upon the ecological and environmental contexts within the oral niche (4–7). For instance, *Streptococcus mutans*, a key contributor to dental caries, exerts its pathogenic potential primarily under conditions of frequent dietary sugar intake, which leads to acid production and subsequent enamel demineralization (8).

In contrast, commensal species are generally regarded as harmless, and in many cases, beneficial to the host. These organisms contribute to oral health through mechanisms such as colonization resistance, where a stable and diverse microbiota prevents the establishment of invading pathogens via nutrient competition, physical exclusion, and production of antimicrobial compounds (7). The classification of a species as a commensal is context dependent. A prime example is *Streptococcus sanguinis*, which is typically abundant in health-associated oral biofilms where it can inhibit *S. mutans* and other pathobionts through hydrogen peroxide production. However, when it gains access to the bloodstream, *S. sanguinis* can cause serious extraoral infections, such as infective endocarditis or, more rarely, brain abscesses in immunocompromised individuals (9).

Despite their dominance in the health-associated oral biofilm and their ecological importance, oral commensals remain underexplored compared to pathobionts and other human associated pathogens (7). The molecular strategies employed by commensals to maintain microbial homeostasis, a concept we define as molecular commensalism (7, 10, 11), have been most extensively studied in oral streptococci. Recent investigations have highlighted the cooperative and spatial interactions between streptococci and *Corynebacterium* spp., two of the most abundant genera in health-associated oral biofilms (11–15). Notably, *Corynebacterium spp.*, long known primarily for pathogenic species such as *C. diphtheriae*, have gained attention for its role as a commensal in the oral cavity as well as the nasopharynx (11, 16, 17). For example, *Corynebacterium* spp. form striking “corncob” structures with streptococci *in vivo*, in which a central *Corynebacterium* rod is surrounded by chains of cocci as demonstrated with fluorescence *in situ* hybridization (FISH) using native human biofilm samples (18). Coculture experiments investigating the corynebacterial-streptococcal interaction *in vitro* have shown that *C. durum* induces up to 10-fold elongation of *S. sanguinis* chains. This effect persists even in transwell setups, indicating that direct contact is not required, instead a diffusible factor mediates the response (15). The study also identified specific fatty acid cargo within *Corynebacterium* extracellular membrane vesicles (EMVs) as the responsible agents for chain elongation. These findings suggest a novel and perhaps widespread form of interspecies communication in the oral biofilm via diffusible EMVs.

The eHOMD lists 20 *Corynebacterium* species inhabiting oral sites, with *C. durum* and *C. matruchotii* among the most abundant (18, 19). While recent studies have begun to explore their roles in oral microbial ecology, information about phenotypes and genomic arrangements as well as cultured representatives remain limited. To address this gap, we have isolated and characterized new oral *Corynebacterium* strains. Here, we present their phenotypic properties, compare their genomes to published strains and discuss their potential ecological roles within the oral microbiome.

## Materials and Methods

### Bacterial strains and media

Strains are described in Supplemental Table 1. Media used for bacterial growth includes Brain Heart Infusion (BHI; Becton Dickinson & Co., MD, USA) and Artificial Saliva Medium (ASM) (20). Bacteria were grown in an ambient atmosphere chamber with 5% CO_2_ at 37°C unless otherwise specified. Strain isolation: Corynebacterial strains were isolated as previously described (15, 21). Briefly, after saliva collection from volunteers, samples were placed on ice to rest prior to being centrifuged (10 minutes, 4000 rpm, A-4-62 Rotor; Eppendorf 5425 Centrifuge) at 4°C. The pellets were then plated on a selective medium for oral *Corynebacterium* species (OCM). BHI was used as OCM base medium supplemented with galactose and bromocresol purple to distinguish *C. durum* from *C. matruchotii*. *C. matruchotii* is unable to catabolize galactose (22). Two antibiotics, fosfomycin and amphotericin B, were included to inhibit the growth of other oral microbes, in particular, oral streptococci and fungi. Two new isolates were chosen for comprehensive characterizations. *C. durum* JJ2, and *C. argentoratense.* The protocol for corynebacterial isolation was approved by the OHSU institutional IRB, study number STUDY00016426.

### Transwell Assay

To test if the newly isolated corynebacterial strains induce chain elongation in *S. sanguinis* as reported before (15), transwell assays were performed. Briefly, fresh overnight cultures of bacterial cells were inoculated in the upper (*C. durum* JJ2, *C. argentoratense)* and lower (*S. sanguinis* SK36) chambers of the well and incubated overnight at 37 °C in a 5% CO_2_ atmosphere. The BHI medium for growth was shared between both bacterial strains, while the cells could not pass the 0.4 μm membrane barrier of the transwell inserts (Transwell^®^ Clear Inserts, Corning, AZ, USA). Cell morphology was then examined in comparison to SK36 in the BHI medium control using an Olympus IX73 inverted microscope. Images were acquired using imaging software platform cellSens (Olympus) to document the elongation phenotype.

### Autoaggregation Assay

An autoaggregation protocol was adapted from Nwoko et al. (23) Briefly, bacterial cells were grown aerobically (5% CO_2_) at 37°C in BHI media for 16 hours. Overnight cultures (OD_600_ > 1.00) were centrifuged (10 minutes, 3000 rpm, A-4-62 Rotor; Eppendorf 5425 Centrifuge) and the medium was removed. Next, the pellet was resuspended in 1 mL aggregation buffer (1mM Tris, 2mM CaCl_2_, 3mM MgCl_2_, 150mM NaCl, buffered to pH 7.4 with HCl) and centrifuged again. The buffer was removed, and cells were resuspended in 1 mL aggregation buffer and diluted to OD_600_ ∼ 1.0 in 1 mL aggregation buffer in polyurethane cuvettes. Samples were incubated at room temperature for 120 minutes and the OD_600_ was measured in increasing intervals up to 120 minutes using a BioPhotometer Plus UV/Vis Photometer (Eppendorf, Germany).

### Crystal Violet biofilm assay

Bacteria were grown overnight in BHI and diluted to OD_600_ = 0.1 in 5 mL BHI. Using a 24 well plate, 500 µL of each culture was added to each well and left to grow for two days. Media was removed and washed 2x with sterile dH_2_O. 600 µL of 0.01% Crystal Violet (CV) was added to each well and left for 30 minutes to stain while rocking on a plate shaker. Next, the CV solution was removed and washed once with dH_2_O and left to dry. To solubilize, 1 mL of 70% ethanol was added to each well and left to sit for 30 minutes while rocking. The absorbance of each well was read at OD_590_ using BioTek Cytation 5 (Agilent, Santa Clara, CA).

### Evaluation of oxygen tension dependent growth conditions

To determine growth under different oxygen tensions, cultures were grown overnight in BHI and diluted to an OD_600_ of 0.1. Next, the cultures were spotted in a 10-fold serial dilution on BHI agar plates and incubated at 37°C either in an ambient atmosphere chamber with 5% CO_2_ or an anaerobic chamber containing an atmosphere of 90% N_2_, 5% CO_2_, and 5% H_2_. Bacterial growth was assessed after 2 days of incubation and photographed for documentation.

### Scanning Electron Microscopy

Cultures were grown in ASM supplemented with glucose, fructose, or mucin, at 1% w/v, and diluted to OD_600_ = 0.3 in ASM. 1 mL of culture was added to 13 mm Thermanox discs situated in a 24 well plate. Biofilms were grown overnight at 37°C with 5% CO_2_ supplementation. The media was removed, and biofilms were fixed with 2% glutaraldehyde in Sorensen’s buffer for 24 hours at 4°C. Biofilms were prepared and imaged at the OHSU Multiscale Microscopy Core, a member of the OHSU University Shared Resource Cores (RRID:SCR_009969). Samples were sputter coated with 10-nm thick carbon (ACE600 coater). Imaging was then performed using a Helios Nanolab 660 dual-beam scanning electron microscope (FEI).

### Confocal Laser Scanning Microscopy (CLSM)

Cultures were grown in BHI and diluted to 0.1 in fresh BHI. 700 µL of diluted culture was added to ibidi^®^ µ-slide 4 well chambered coverslips. After 1 day of growth, cultures were supplemented with 5 µL of 5 mM Syto 9 stain solution to result in a 0.0357 mM solution and taken to the Advanced Light Microscopy Core at Oregon Health Science University for microscopy. Subsequent images were analyzed using Imaris 3D rendering with Blend and MIP (Maximum Intensity Progression) thumbnails.

### Corynebacterial API test

Newly isolated strains were characterized using the commercial API^®^ Coryne system (BioMérieux, France). In brief, this biochemical reaction system consists of dehydrated substrates for 11 enzymatic activities (nitrate reductase, pyrazinamidase pyrrolidonyl arylamidase, alkaline phosphatase, glucuronidase, β-galactosidase, α-glucosidase, *N*-acetyl-β-glucosaminidase, glucosaminidase, esculin, urease, and hydrolysis of gelatin) and eight sugar fermentations (glucose, ribose, xylose, mannitol, maltose, lactose, sucrose, and glycogen). Bacterial samples were prepared and transferred to API^®^ Coryne strips according to the manufacturer’s instructions. The strips were then incubated at 37°C for 24 hours. The readings, except for the esculin, urease, and gelatin tests, were performed after adding the appropriate reagents. The fermentation reactions were considered positive when they turned yellow. Identification and interpretation were conducted according to the manufacturer’s instructions. Catalase activity was determined by adding a drop of hydrogen peroxide (3%) to the esculin test after 24h.

### Extracellular membrane vesicle (EMV) concentration

EMV isolation and characterization followed a previously published protocol (24). Bacterial cells were grown in 25 mL ASM overnight and diluted to 1.0 in 250 mL ASM media supplemented with glucose or sucrose at 0.6% as relevant carbohydrates known to influence the oral biofilm. Cultures were left to grow overnight on a shaking incubator at 37°C and 150 rpm. Samples were centrifuged (15 minutes, 3750 rpm, A-4-62 Rotor; Eppendorf 5425 Centrifuge) at 4°C. The resulting supernatant was filter sterilized with a 0.45 µm pore and then divided amongst Vivaspin 20 ultracentrifugation units (100 kDa MWCO, GE Healthcare) columns for further concentration. The columns were centrifuged at 4°C at increasing times of 15-45 minutes and 4000 rpm (A-4-62 Rotor; Eppendorf 5425 Centrifuge). Once the supernatant was concentrated to < 5mL, the resulting solution was ultracentrifuged for 2 hours at 37,000 rpm and 4°C (Beckman Coulter, Optima XL-100K Ultracentrifuge; rotor type 50.3TI). The final precipitate was resuspended in 1 mL PBS and stored at -80°C for long term storage. EMVs were quantified with NTA using ZetaView (Particle Metrix, Germany), scanning 11 cell positions with 60 frames per position for every measurement. These positions were analyzed by ZetaView software version 8.05.12 with the following parameters: laser wavelength (488 nm), filter wavelength (scatter), maximum particle size (1,000), minimum particle size (10), minimum particle brightness (20).

### Corynebacterium pangenome and phylogenomic tree

To compare the pangenome and phylogeny of the strains examined here, Anvi’o (development version) was used to generate a pangenome of the 4 strains examined here, plus the additional *C. matruchotii* reference strain NCTC10206. All files and code used here are available at https://github.com/jonbakerlab/Corynebacterium-pangenome. The pangenome was used to select 14 single-copy core amino acid sequences with maximum sequence heterogeneity (but no gaps in the alignment) with which to perform phylogenomic analysis, also performed using Anvi’o. A *C. glutamicum* genome was added to the phylogenomic analysis as a non-oral outgroup, however the analysis unexpectedly showed that *C. glutamicum* was more closely related to *C. matruchotii* than the other strains. NCBI accession numbers are: *C. durum* JJ2: CP198957; *C. durum* JJ1: CP199749; *C. argentoratense* MB1: CP199748

## Results

### Isolation of oral corynebacterial species

Using an established protocol for the selective isolation of oral *Corynebacterium* species (15, 21), six putative isolates were obtained. 16S rRNA gene sequencing identified one isolate as *C. durum* with 99.25% sequence identity based on NCBI BLASTn analysis (standard database, default parameters). The remaining five isolates were identified as *C. argentoratense*, ranging between 96–99% sequence identity (data not shown). For subsequent characterization, the *C. durum* isolate designated as strain JJ2 and one *C. argentoratense* isolate designated as strain MB1 were selected.

### Induction of Cell Chain Elongation in *S. sanguinis*

One of the few well-characterized *in vitro* interspecies interactions involving oral *Corynebacterium* species is the induction of cell chain elongation in the commensal *S. sanguinis*, mediated by extracellular membrane vesicles (EMVs) (15). This phenotype appears to be specific to *S. sanguinis* and has not been observed in other tested oral streptococci. The effect has previously been demonstrated for both *C. durum* and *C. matruchotii* (15). To determine whether the newly isolated strains also elicit this response, transwell assays were conducted with *S. sanguinis* SK36 and physically separated from corynebacterial species via transwell inserts. Consistent with previous findings, both *C. durum* JJ2 and *C. argentoratense* MB1 induced marked chain elongation in *S. sanguinis* following overnight incubation (Fig. 1A). Of note, all newly *C. argentoratense* strains isolated induced cell chain elongation (Supplemental Fig. 1). These results suggest that the new isolates likely produce EMVs capable of diffusing through the transwell membrane to trigger the observed phenotypic change in *S. sanguinis* chain formation.

**Figure 1:**
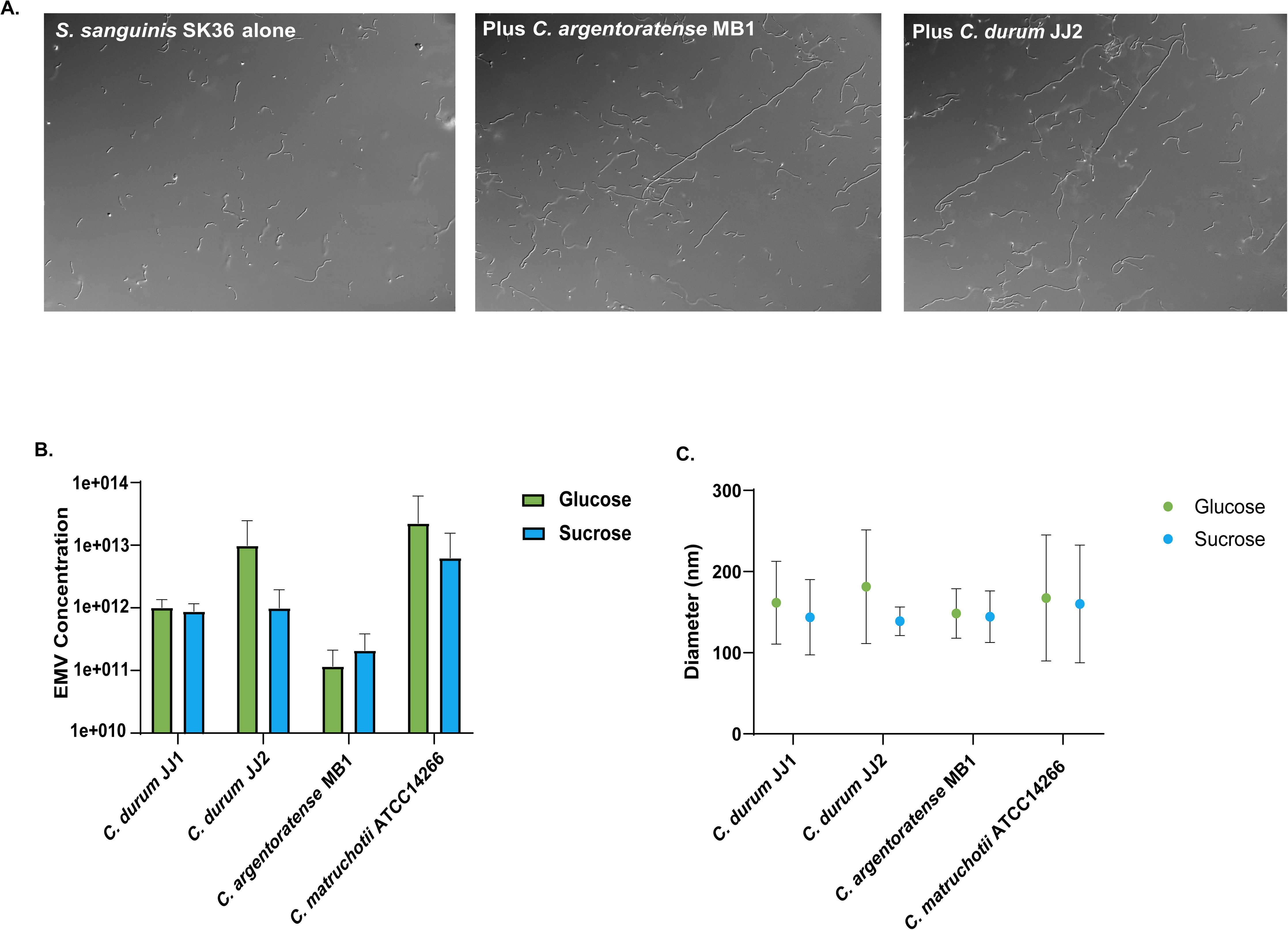
A. Transwell assays for *S. sanguinis* SK36 alone, SK36 with *Corynebacterium argentoratense* MB1, SK36 with *Corynebacterium durum*. B. Bar graph showing the concentration of EMVs produced by *Corynebacterium* species grown in Artificial Saliva Media with either glucose or sucrose supplemented (n=3, standard deviation). C. Graph showing the varying diameter of EMVs produced by *Corynebacterium* species grown in Artificial Saliva Media with either glucose or sucrose supplemented. (n=3, standard deviation)

### Characterization of Corynebacterial EMV Production

Extracellular membrane vesicles play a critical role in mediating interactions between oral *Corynebacterium* species and other members of the oral microbiome. Notably, EMVs have been shown to induce chain elongation in several isolates of *S. sanguinis* and to inhibit hyphal formation in *Candida albicans* (15, 25). To further investigate if relevant ecological factors implicated in caries development are influencing EMV production in *Corynebacterium*, we examined the impact of glucose and sucrose supplementation on EMV biogenesis. All four tested *Corynebacterium* strains produced EMVs when cultured in ASM supplemented with either glucose or sucrose. EMV diameters and concentrations were comparable between the two carbohydrate conditions (Fig. 1B and C), suggesting that the type of sugar does not significantly influence vesicle production. Among the carbohydrate conditions, *C. durum* JJ2 produced the largest vesicles in glucose, and *C. matruchotii* produced the largest vesicles in sucrose. *C. matruchotii* yielded the highest EMV concentrations reaching up to 10¹³ particles per mL, which correlates with its enhanced growth and higher final cell density following overnight incubation. *C. durum* JJ1 and *C. argentoratense* MB1 yielded lower EMV concentrations and diameters across both carbohydrate conditions. Visible differences in EMV diameter are seen for *C. durum* JJ2 in glucose in contrast to sucrose, yet the data is not significant (Fig. 1B and C). Overall, all four *Corynebacterium* strains produced substantial amounts of EMVs under both carbohydrate conditions, and EMV production appeared largely unaffected by the specific sugar source.

### Comparison of Corynebacterium autoaggregation

Bacterial aggregation and autoaggregation are commonly mediated by adhesins, including fimbriae or pili, as well as specific surface proteins as demonstrated for the streptococcal antigen I/II family and CshA/B (26, 27). In addition, secreted factors like polysaccharides and extracellular DNA (eDNA) can contribute to aggregation behavior (28). However, in oral *Corynebacterium* species, no specific proteins or molecular factors have yet been identified that mediate aggregation or autoaggregation. To assess their potential for autoaggregation, we compared the autoaggregation behavior of all four oral *Corynebacterium* isolates (Fig. 2A). Both *C. durum* strains exhibited rapid autoaggregation, with visible clumping occurring within 15 minutes of incubation in aggregation buffer (Fig. 2B). In contrast, *C. matruchotii* and *C. argentoratense* showed minimal aggregation, even after 60–90 minutes. *S. sanguinis* was included as a negative control and did not aggregate during the 120-minute observation period. Notably, the time-dependent autoaggregation rate of *C. durum* JJ1 and JJ2 was approximately fivefold higher than that of the other two *Corynebacterium* species (Fig. 2A).

**Figure 2:**
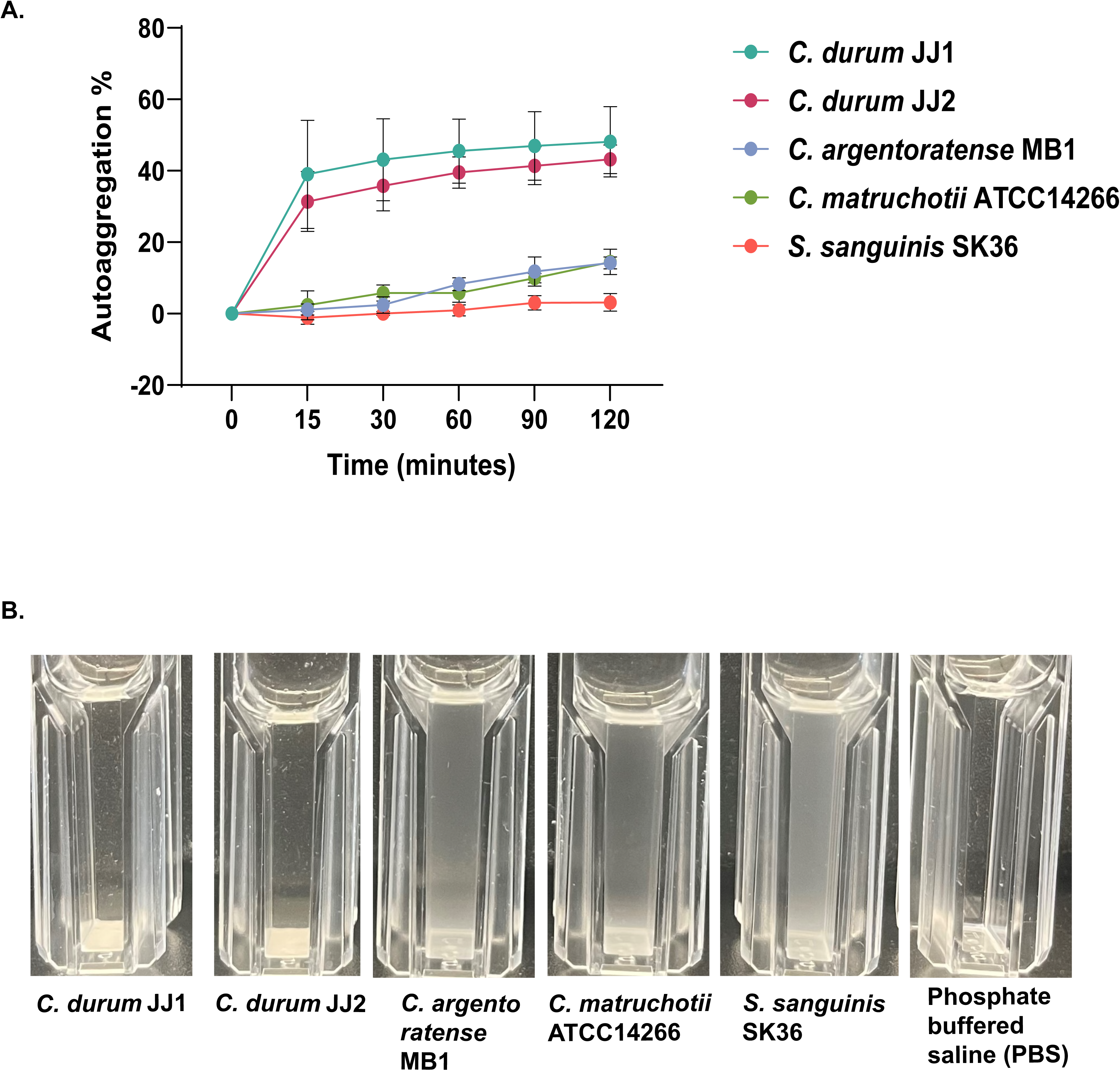
A. Average autoaggregation percentage of *Corynebacterium* species after 120 minutes of static incubation (n=3, standard deviation). B. Bacterial species after 120 minutes of static incubation. 1 = *C. durum* JJ1, 2 = *C. durum* JJ2, 3 = *C. argentoratense*, 4 = *C. matruchotii* ATCC, 5 = *S. sanguinis* SK36, 6 = PBS control (representative picture of n=3).

### Biofilm Quantification

Bacterial aggregation is a key mechanism that can directly impact biofilm formation (23). To assess biofilm production among the corynebacterial strains, we employed the standard crystal violet staining assay. As shown in Figure 3A, both *C. durum* strains retained more crystal violet compared to the other species, indicating the formation of thicker biofilms. In contrast, *C. argentoratense* MB1 and *C. matruchotii* retained less crystal violet, suggesting the formation of thinner biofilms. The observed differences in autoaggregation, with strains JJ1 and JJ2 exhibiting enhanced aggregation capacity, along with their increased biofilm formation, suggest a possible correlation between these two processes.

**Figure 3:**
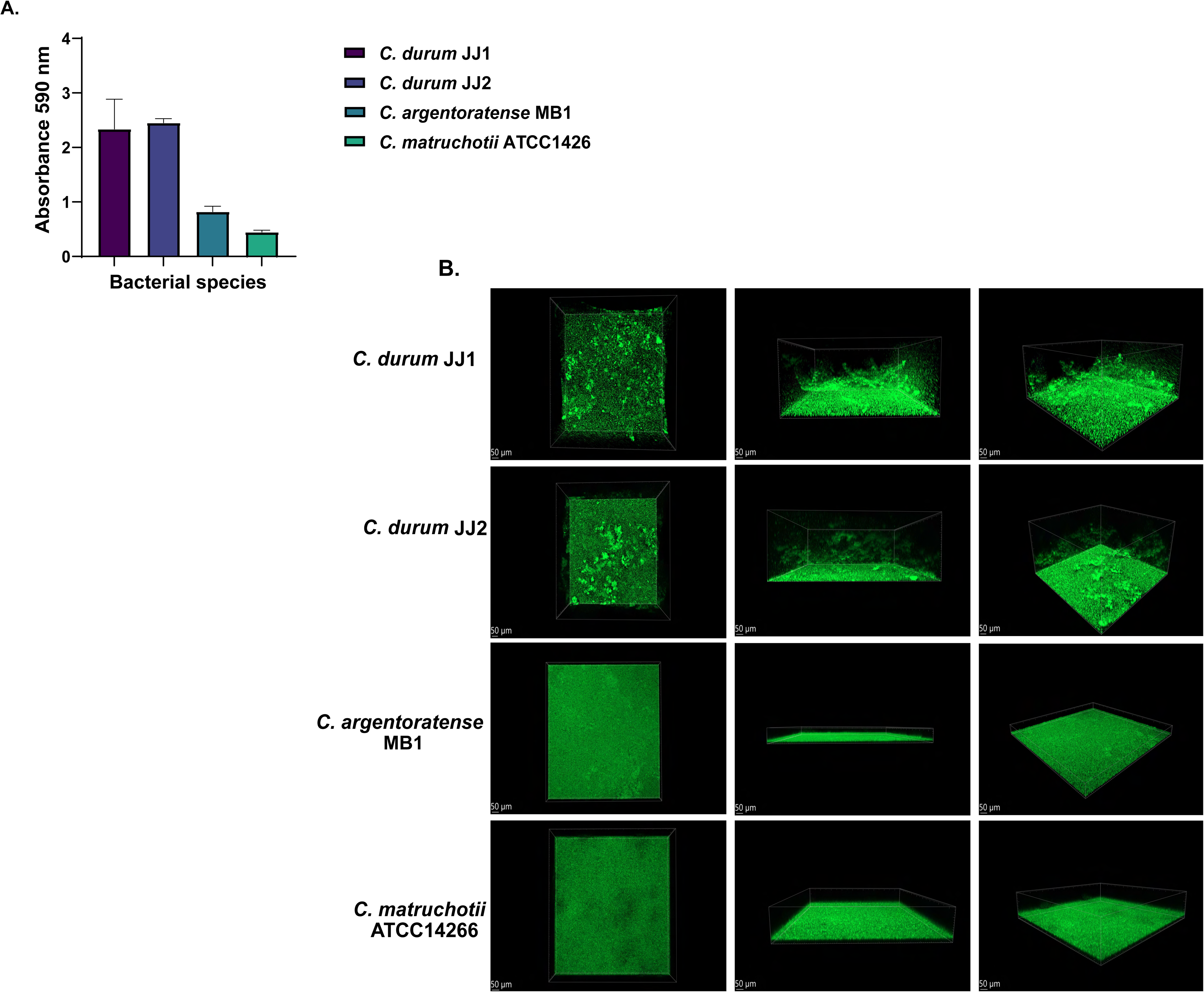
A. Average absorbance of crystal violet-stained biofilms read at absorbance 590 nm. (n=3, standard deviation) B. Confocal laser scanning microscopy (CLSM) of the four *Corynebacterium* strains. 3D rendering performed using Imaris with a maximum intensity progression performed.

### CLSM Imaging of 3D Biofilm Structure

Although crystal violet staining enables a general quantification of biofilm biomass, it does not provide information regarding the structural organization or architecture of the biofilm. To assess the three-dimensional structure of the biofilms, Confocal Laser Scanning Microscopy (CLSM) was performed on all four isolates (Fig 3B). Consistent with the crystal violet assay, CLSM revealed striking differences in biofilm morphology: the *C. durum* strains formed thick, highly aggregated biofilms, whereas *C. argentoratense* MB1 and *C. matruchotii* produced thin, uniform biofilm mats. These findings indicate a consistent phenotypic similarity between the two *C. durum* strains across multiple assays, crystal violet staining, autoaggregation, and CLSM highlighting their distinct ability to form robust, structured single-species biofilms compared to *C. argentoratense* MB1 and *C. matruchotii*.

### SEM Analysis of Biofilms and EPS Structures

We previously demonstrated using SEM that *C. durum* produces an elaborate network of extracellular polymeric substances (EPS), enmeshing cells within a dense biofilm matrix (13). To further investigate the extracellular matrix and cellular morphology of the four *Corynebacterium* strains, SEM imaging was performed on cells grown in ASM supplemented with different carbohydrate sources: glucose, fructose, and mucin. As shown in Fig. 4, both *C. durum* strains exhibited prominent extracellular matrix structures, particularly in the presence of glucose, where a dense, highly branched EPS network surrounded the cells. This network was less developed when cells were grown with fructose. Interestingly, mucin supplementation also led to the formation of extracellular material, though it appeared morphologically distinct. Notably, both *C. durum* strains (JJ1 and JJ2) showed similar cellular morphology and EPS features under all tested conditions. In contrast, *C. argentoratense* MB1 and *C. matruchotii* did not form extensive EPS networks in the presence of glucose or fructose. However, mucin-supplemented conditions induced the production of some extracellular material in these species, albeit to a lesser extent than in *C. durum*. Additionally, marked differences in cell morphology were observed: *C. matruchotii* formed elongated cells, while *C. argentoratense* exhibited much shorter, stubbier cells compared to the other three strains (Fig 4). Together, these findings highlight distinct structural phenotypes among oral *Corynebacterium* species, with *C. durum* demonstrating a unique capacity to produce a dense EPS matrix and maintain consistent cell morphology across nutrient conditions, supporting its role in robust biofilm formation.

**Figure 4:**
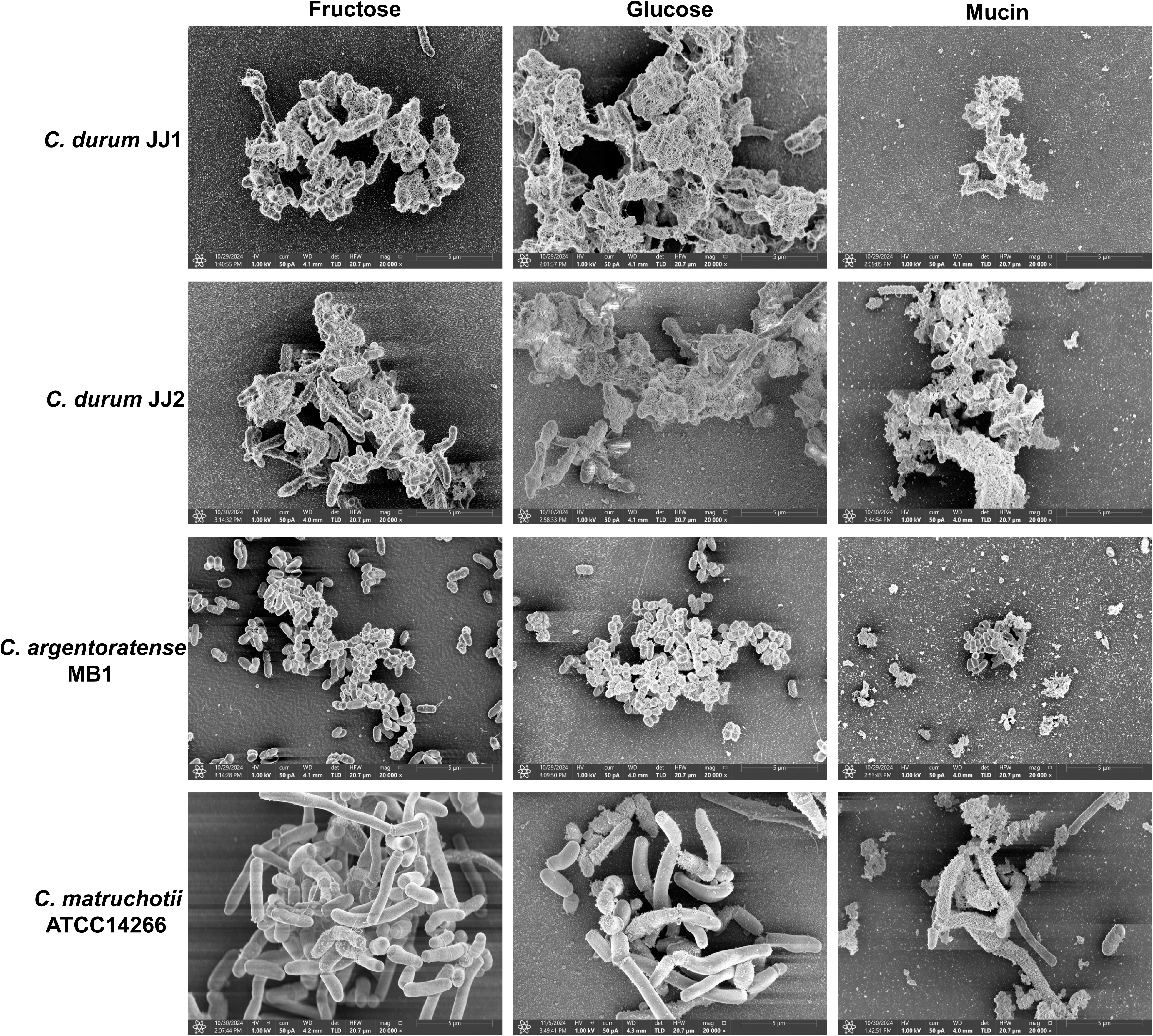
Scanning Electron Microscopy (SEM) of four *Corynebacterium* isolates with fructose, glucose, and mucin supplemented growth media at 20,000x magnification.

### Biochemical Analyses of New Isolates

To further characterize the recently isolated oral *Corynebacterium* strains, biochemical profiling was performed using the API® Coryne system. Both *C. durum* isolates (JJ1 and JJ2) displayed largely similar biochemical profiles, with minor differences observed in the β-galactosidase activity (β-GAL) with lower activity of JJ1 compared to JJ2 and maltose (MAL) fermentation tests, which seems to be absent in JJ1 (Supplemental Fig. 2). In contrast, *C. argentoratense* MB1 showed a distinct biochemical signature, lacking positive reactions for nitrate reduction (NIT), and esculin hydrolysis (ESC), and only seemed to be able to ferment glucose (GLU) under the test conditions. Additionally, *C. matruchotii* differed by exhibiting positive reactions for pyrrolidonyl arylamidase (PyrA) and α-glucosidase (α-GLU), which were negative in the other isolates (Supplemental Fig. 2). Overall, the majority of test results were consistent across all four strains, with only a subset showing species-specific differences. These findings agree with previous biochemical profiling conducted by our group (29), where *C. durum* strain JJ1 and *C. matruchotii* ATCC displayed similar metabolic patterns.

### The effect of oxygen tension on growth

A notable difference in the biochemical profiles of the four *Corynebacterium* strains was the absence of nitrate reduction in *C. argentoratense* MB1. While corynebacteria are generally considered as aerobic bacteria, some species are facultative anaerobes, such as *C. glutamicum*, which is capable of anaerobic growth through nitrate respiration and fermentation, resulting in mixed acid production (30). In the oral cavity, corynebacteria encounter oxygen-rich conditions during early biofilm formation, but may experience oxygen limitation in more mature, stratified biofilms (31, 32). To assess their growth potential under varying oxygen levels, all four strains were cultured under aerobic (ambient air with 5% CO_2_) and anaerobic (90% N^2^, 5% CO^2^, 5% H^2^) conditions (Fig. 5). BHI agar plates spotted with *Corynebacterium* cultures were incubated at 37°C for 48 hours. Under aerobic conditions, all four strains displayed robust and consistent growth. In contrast, anaerobic conditions led to a general reduction in growth, with *C. argentoratense* MB1 showing no detectable growth. These findings suggest that *C. argentoratense* may have limited adaptive capacity for survival in anoxic environments, potentially influencing its spatial niche within the oral biofilm.

**Figure 5.**
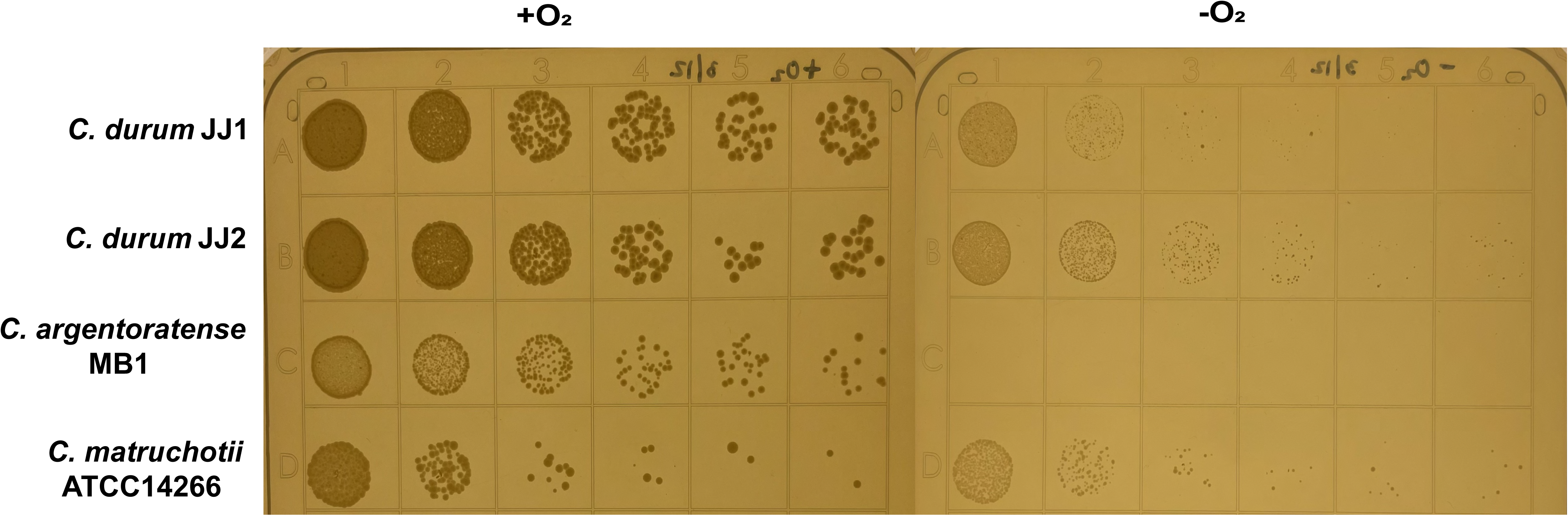
BHI agar plates with spotted serially diluted *Corynebacterium* species grown for 2 days in either aerobic (+O_2_) or anaerobic (-O_2_) chambers.

### Comparative Genome Analysis

The general genome feature analysis showed that the GC content was similar across species (∼57%), while genome size and total gene count were comparable between *C. matruchotii* and *C. durum* but markedly reduced in *C. argentoratense* MB1 (Fig 6A). Pangenomic and phylogenomic analyses were performed to compare the genomes of *Corynebacterium* isolates. For comparison, *C. glutamicum*, a prominent non-oral Corynebacterium used for industrial large scale amino acid production, was also included. Phylogenomic analyses indicated that *C. glutamicum* was more closely related to *C. matruchotii*, while *C. durum* was more distant and *C. argentoratense* MB1 was most distantly related. (Fig 6B). In line with the phylogenomic data, pangenomic analysis of the newly isolated and sequenced *C. durum* JJ2 and *C. argentoratense* with the published sequences of *C. durum* JJ1 and two *C. matruchotii* strains revealed considerable pangenomic overlap within *C. durum* and *C. matruchotii*, but a striking divergence from *C. argentoratense,* which had 807 unique genes. (Fig. 6C). Analysis of shared genes further supported this divergence: *C. matruchotii* and *C. durum* collectively shared 240 clusters, whereas *C. argentoratense* shared only 107 with *C. durum* and 29 with *C. matruchotii*. Across all five genomes, 910 core genes were shared out of 11,645 total (Fig. 7A). COG20 functional categorization indicated that the “Translation, ribosomal structure and biogenesis” category was the most abundant in all genomes, with 1.5–2× more hits than other categories (Fig 7B). Targeted examination of nitrate reduction genes revealed that *C. durum* and *C. matruchotii* harbor three of the four *narGHIJ* operon genes (*narG*, *narI*, *narJ*) essential for nitrate reduction and anaerobiosis in *E. coli* and other bacteria. These genes were absent in *C. argentoratense* MB1, consistent with its lack of nitrate reductase activity and inability to grow under anaerobic conditions (Fig. 7C). In summary, *C. argentoratense* exhibits substantial genomic divergence from *C. durum* and *C. matruchotii*, reflecting distinct functional capacities and potential adaptation to unique oral ecological niches.

**Figure 6:**
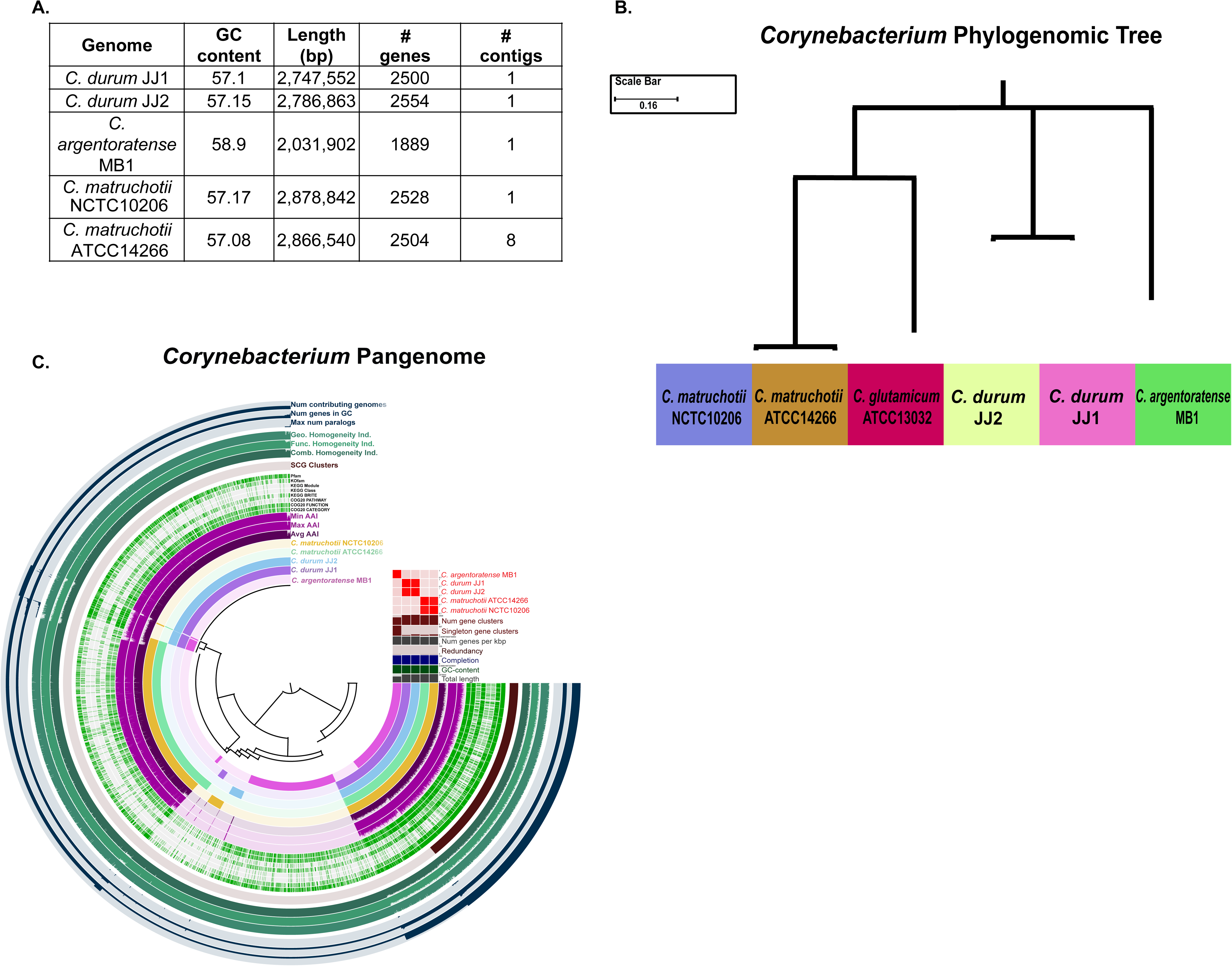
A. Chart displaying unique characteristics to each genome. B. Phylogenomic Tree of 6 Corynebacterial genomes, specifically the evolutionary differences between the shared single copy core genes present within all genomes. C. Pangenome assembly of 5 Corynebacterium genomes using bioinformatics tool Anvio.

**Figure 7:**
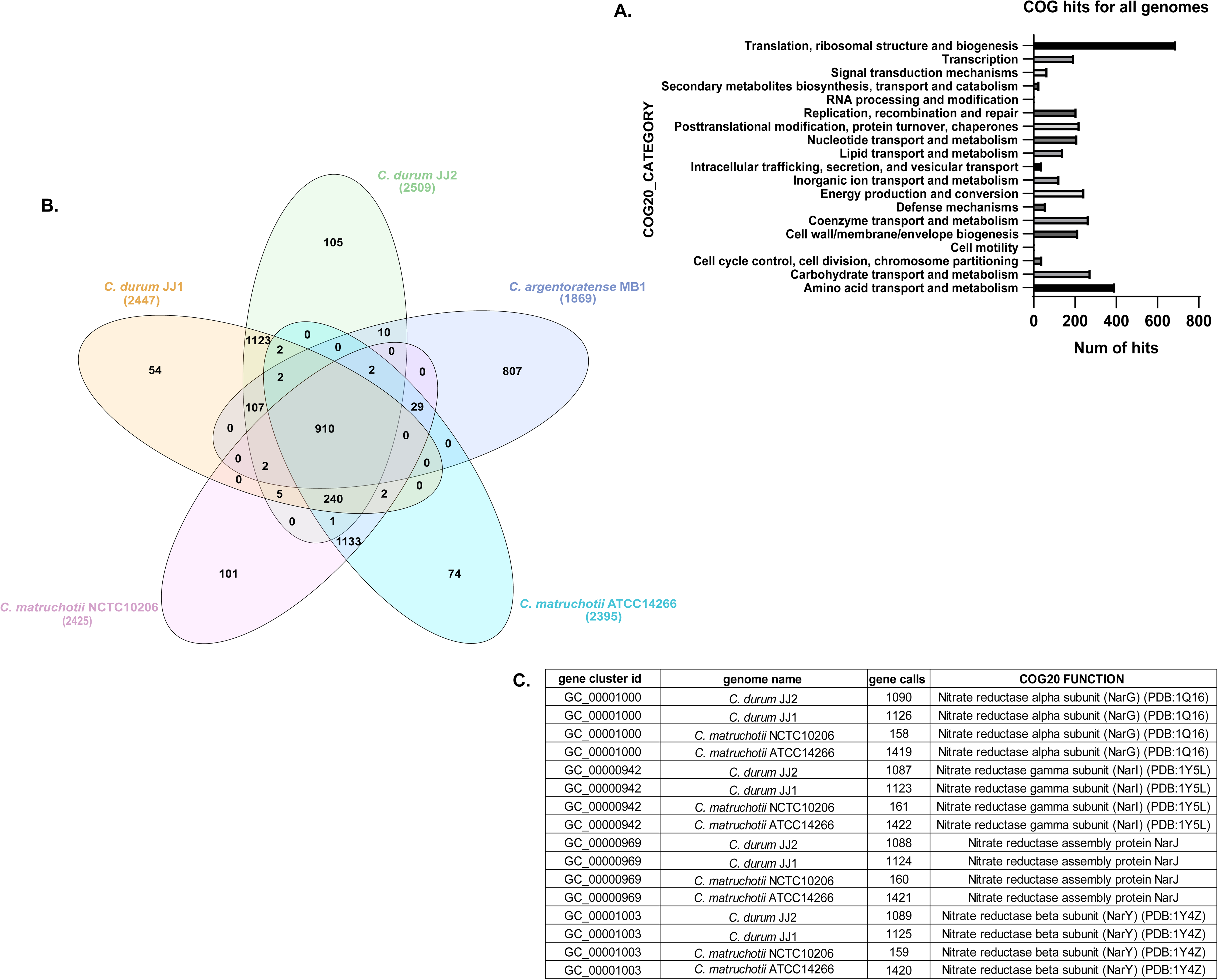
A. Venn diagram showing the gene clusters in the five genomes used for the pangenome. B. Number of hits for each COG category within the pangenome. C. Chart showing unique nitrate genes to four out of five genomes (no homologs present in *C. argentoratense*).

## Discussion

Oral microbiology has traditionally focused on a limited number of bacterial species strongly associated with oral diseases, for example, *S. mutans* in dental caries and *P. gingivalis* and *Treponema denticola* in periodontitis (33–36). These studies have greatly advanced our understanding of specific pathogenic mechanisms and contributed to the broader field of microbial virulence. However, despite these insights, the overall prevalence of caries has only slightly improved and periodontal disease has remained largely unchanged (37). This paradox reflects a shift in our understanding of oral disease etiology. It is now widely accepted that diseases such as caries and periodontitis are polymicrobial in nature, resisting traditional pathogen-centered diagnostic and therapeutic approaches (4, 38). The concept of microbial dysbiosis has emerged to explain how a once-balanced microbial community can shift toward a disease-associated state. Enabled by high-throughput sequencing technologies, studies of the oral microbiome have revealed substantial compositional and functional changes in microbial communities associated with disease development (39–41). Crucially, these changes reflect a transition from a healthy (eubiotic) microbiome to a dysbiotic one and not through the invasion of new pathogens, but through behavioral and metabolic shifts in resident species that adapt to the altered oral environment (6). This paradigm shift has underscored the need for a more holistic understanding of oral disease, one that accounts not only for pathogenic processes but also for the ecology and function of the healthy microbiome. In this context, oral commensals, particularly commensal streptococci, have received renewed attention (7). These organisms contribute to oral health through a variety of mechanisms that enhance their fitness and confer colonization resistance. One such mechanism is hydrogen peroxide (H^2^O^2^) production via the SpxB pathway, which inhibits the growth of H^2^O^2^-sensitive pathobionts such as *S. mutans* and *P. gingivalis* (42).

Interest in other abundant yet understudied members of the oral microbiome has also grown. A landmark study using fluorescence in situ hybridization (FISH) to characterize the spatial organization of oral biofilms highlighted the high abundance and central positioning of corynebacteria within supragingival biofilms (18). This finding catalyzed interest in the ecological role of oral *Corynebacterium* species, which, despite their prevalence, had long remained functionally obscure (43). This renewed focus coincides with broader interest in corynebacteria colonizing other mucosal surfaces, such as the nasopharynx (16) and the ocular surface (44), suggesting that these bacteria may play important and possibly conserved roles across different host-associated microbial communities. The corynebacterial species examined here, a newly isolated *C. durum* strain and the less commonly studied *C. argentoratense,* expand our understanding of phenotypic traits within this underappreciated genus.

Overall, their genome features are consistent with other human-associated corynebacteria. The G+C content of *C. durum* (JJ1 and JJ2) and *C. matruchotii* is ∼57%, whereas *C. argentoratense* MB1 is slightly higher at 58.9%, matching previously reported values for three *C. argentoratense* strains isolated from the respiratory tract and from blood. In sum, their G+C content falls within the published range for human-associated corynebacteria (approximately 53–58%) (22, 45). Similarly, the genome sizes of *C. durum* and *C. matruchotii* fall within the expected range for corynebacteria, averaging around 3 Mb. In contrast, *C. argentoratense* MB1 is considerably smaller, at just over 2 Mb. This reduction is also evident in its gene content, with only 1,889 predicted genes compared to approximately 2,500 in the other two species (45, 46).

The phenotypic differences observed across the range of experiments highlight a strong similarity between the two *C. durum* strains JJ1 (previous and published isolate) and JJ2, which is to be expected, while also revealing partial phenotypic overlap between *C. argentoratense* MB1 and *C. matruchotii*. This is particularly evident in biofilm formation and autoaggregation assays, which indicate that certain characteristics are conserved between *C. argentoratense* and *C. matruchotii*. However, phylogenomic tree analysis reveals a different evolutionary relationship: *C. durum*, *C. matruchotii, C. argentoratense* and *C. glutamicum* (used as non-oral *Corynebacterium* for comparison) share 14 core genes, yet only *C. matruchotii* and *C. glutamicum* cluster on the same branch, suggesting that they come from a recent common ancestor, while the other two species have evolved in a different manner (Fig. 6B). These findings may reflect spatial niche differences within the oral and upper respiratory microbiomes. For instance, *C. argentoratense* has been isolated from a variety of anatomical sites including the throat (particularly in patients with tonsillitis), upper respiratory tract, blood cultures, and mucosal biofilms (45, 47). In contrast to *C. durum* and *C. matruchotii*, which are well-represented in eHOMD with 8 and 11 genome entries respectively, *C. argentoratense* is notably absent from the eHOMD genome table. In the present study we isolated *C. argentoratense* from saliva. Despite current insights, the role of *C. argentoratense* in oral ecology remains largely unresolved. Determining whether it is a transient inhabitant or a persistent community member, and whether candidate virulence determinants contribute to disease development, merits further systematic investigation.

One of the most distinguishing biochemical features of *C. argentoratense* MB1 is its inability to reduce nitrate, as demonstrated in API strip assays (Supplemental Fig. 2). The capacity for nitrate reduction, whether to nitrite or further downstream products, is an ecologically relevant trait among oral bacteria (48) and may enable specific taxa to adapt to anaerobic microenvironments. This suggests that *C. argentoratense* may occupy a distinct functional niche within the oral mucosal microbiome that has guaranteed oxygen availability. The well-documented biogeographical distribution of *C. matruchotii* along the supragingival margin, where it serves as a structural scaffold for biofilm development (19, 49), likely selects for its ability to grow even under increasingly anaerobic conditions as the biofilm matures. In addition, *C. matruchotii* has been shown to promote calcification through the deposition of salivary calcium, contributing to dental calculus formation (50), a process that may further limit oxygen diffusion over time. Although the specific spatial distribution of *C. durum* in the oral cavity has not been determined, it is conceivable that, similar to *C. matruchotii*, it is a constituent of the supragingival biofilm.

A notable ecological trait conserved across all oral *Corynebacterium* species examined to date is their ability to induce a species-specific cell chain elongation phenotype in *Streptococcus sanguinis*. This interaction was first studied in detail with *C. durum* and attributed to extracellular membrane vesicles (EMVs), specifically those containing fatty acid cargo (15). *C. durum* secretes fatty acids, most prominently oleic acid, stearic acid, and palmitic acid, and *in vitro* reconstitution of these fatty acids reproduces the chain elongation phenotype in *S. sanguinis* (15). EMV fatty acid content can be modulated by environmental conditions; for instance, growth of *C. durum* in the absence of glucose results in EMVs with a ∼50-fold reduction in fatty acid content, eliminating their chain-elongating activity (15). Although the fatty acid content of EMVs was not determined in the present study, we observed that all four *Corynebacterium* strains produced EMVs of comparable concentration and diameter, suggesting a conserved EMV production capacity responsible for the induction of chain elongation in *S. sanguinis*. Given their ability to diffuse beyond the immediate vicinity of the producing cells, corynebacterial EMVs may influence other microbial community members and even the host. Indeed, EMV uptake by oral epithelial cells has been demonstrated for several oral bacterial species (51–53), underscoring their potential role in modulating both interspecies interactions and host responses within the oral biofilm ecosystem.

A clear distinction was also observed in the metabolic profiles determined using the API Coryne strip test. While not exhaustive, this assay provides a useful means of differentiating corynebacteria and revealed that *C. argentoratense* is unable to metabolize several key carbohydrates that both *C. durum* and *C. matruchotii* can utilize as energy sources. Interestingly, studies on human nasal-associated corynebacteria, including *C. propinquum*, *C. pseudodiphtheriticum*, *C. accolens*, and *C. tuberculostearicum,* have shown that their metabolic pathways are largely conserved, consistent with their shared genomic and pangenomic structures (46). These findings support the conclusion that metabolic capacity is closely tied to niche specialization within the nasopharynx.

By comparison, the results presented here suggest that *C. argentoratense* may be less well adapted to the oral niche, given its reduced metabolic repertoire relative to the other two corynebacterial strains. This raises the possibility that *C. argentoratense* may occur only transiently in the oral cavity. However, it is well established that metabolic interdependencies are widespread within oral biofilms, for example, *Veillonella* spp. relies on lactic acid produced by oral streptococci for growth (54). A similar cross-feeding relationship with other oral bacteria could support the persistence of *C. argentoratense* in this environment. At a minimum, further investigation is needed to clearly define the biogeographical distribution of *C. argentoratense* as well as *C. durum*, whose precise localization within the oral cavity has likewise not been established but is evident for multiple oral microbiome sequencing projects.

In conclusion, our comparative analyses of *C. durum, C. matruchotii* and *C. argentoratense* underscore both the conserved and divergent traits within oral corynebacteria. While *C. durum* and *C. matruchotii* share genomic and phenotypic similarities consistent with stable adaptation to the oral niche, *C. argentoratense* displays reduced genome size, limited metabolic capabilities, and distinct biochemical features that may reflect either niche specialization under restricted conditions or a more transient role in the oral cavity. Its absence from eHOMD despite repeated isolation from oral samples highlights how little is known about its ecological significance. The conserved ability of all tested oral corynebacteria to influence *S. sanguinis* morphology through EMV-associated fatty acids further emphasizes their potential importance as biofilm organizers and modulators of community structure. Collectively, these findings expand our understanding of corynebacterial contributions to oral ecology but also reveal critical gaps, particularly regarding spatial distribution, metabolic dependencies, and potential roles in health and disease, that warrant further investigation.

## Acknowledgement

J.K. acknowledges the support of NIH-NIDCR grants DE029612 and DE029492, J.L.B. of grant DE029228, and J.M. of grant DE028252.

---

**Table 1:** Homology of selected open reading frames.

| *C. argentoratense* | *C. durum* | *C. matruchotti* |
| --- | --- | --- |
| GC00003614<br>Enoyl-CoA hydratase | 30.81% | 27.64% |
| GC00003643<br>Branched chain amino acid efflux pump | 47.22% | 37.02% |
| GC00003689<br>Glutamyl-tRNA reductase | 49% | 43.64% |
| GC00004088<br>Amino acid permease | No significant similarity |  |
| GC00004161<br>Septum formation Initiator FtsB | 40.67% | 36% |
| GC00004206<br>DNA repair protein RecO | 46.18% | 41.86% |
| GC00004250<br>Unknown fnction | 52.43% | No significant similarity |
| GC00004307<br>Maltose 6'-phosphate phosphatase | No significant similarity |  |
| GC00004682<br>N-acetylmuramoyl C-alanine amidase | No significant similarity |  |
| GC00004696<br>Sensor histidine kinase | 50.81% | 45.49% |

## Notes

### Competing Interest Statement

The authors have declared no competing interest.

