## Supplemental Material for "Comparative Characterization Reveals Conserved and Divergent Ecological Traits of Oral Corynebacteria"

| Strains | Characteristics | Reference |
| --- | --- | --- |
| *C. durum* JJ1 | Clinical isolate (GenBank accession number MN251472) | Treerat *et al*. 2020 |
| *C. matruchotii* ATCC 14266 | *C. matruchotii* reference strain | Barrett *et al*. 2001 |
| *C. matruchotii* NCTC 10206 | *C. matruchotii* reference strain #NC_003450 | RefSeq: GCF_900638255.1 |
| *C. durum* JJ2 | Sequenced isolate (GenBank accession number CP198957) | this study |
| *C. argentoratense* MB1 | Sequenced isolate (GenBank accession number) | this study |
| *C. glutamicum* ATCC 13032 | *C. glutamicum* reference strain #NC_003450.3 | RefSeq: GCF_000011325.1 |
| *S. sanguinis* SK36 | *S. sanguinis* wild type | Xu et al. 2007 |

Supplemental Table 1: Bacterial species used in this study.

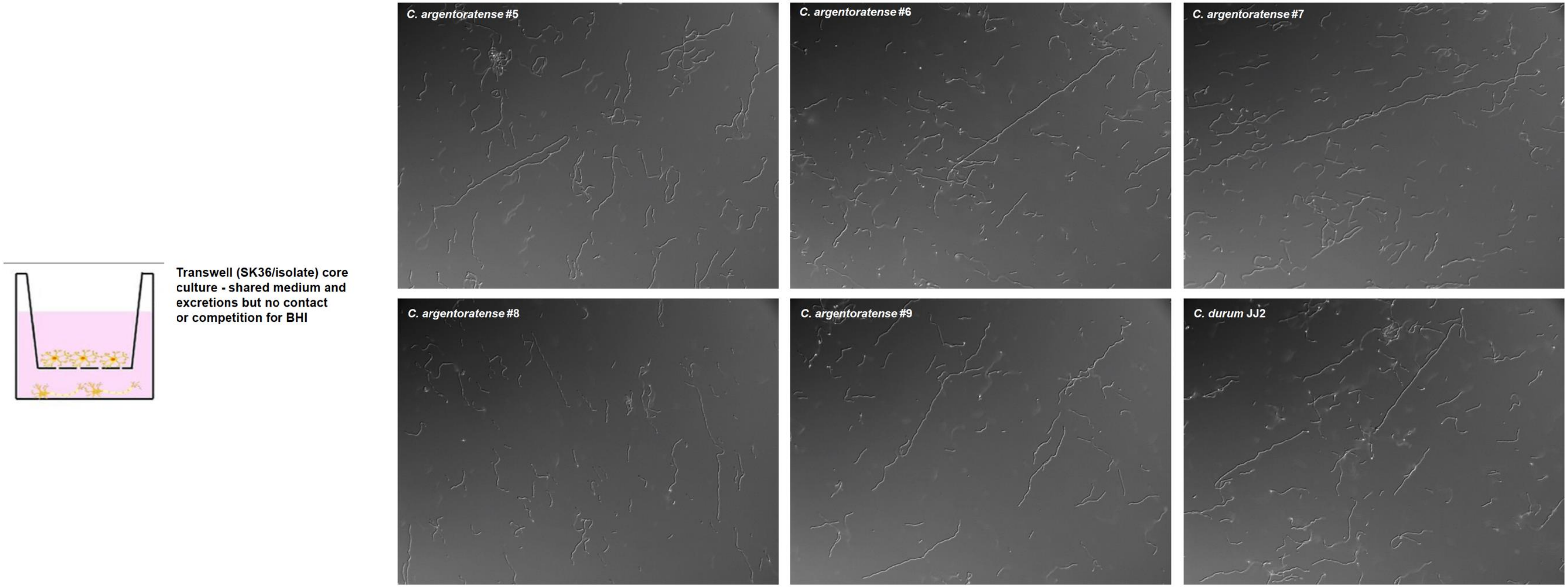


Supplemental Figure 1: Transwell assays for SK36 with *Corynebacterium argentoratense* #5, SK36 with *Corynebacterium argentoratense* MB1, *Corynebacterium argentoratense* #7, *Corynebacterium argentoratense* #8, *Corynebacterium argentoratense* #9, and *Corynebacterium durum* JJ2.

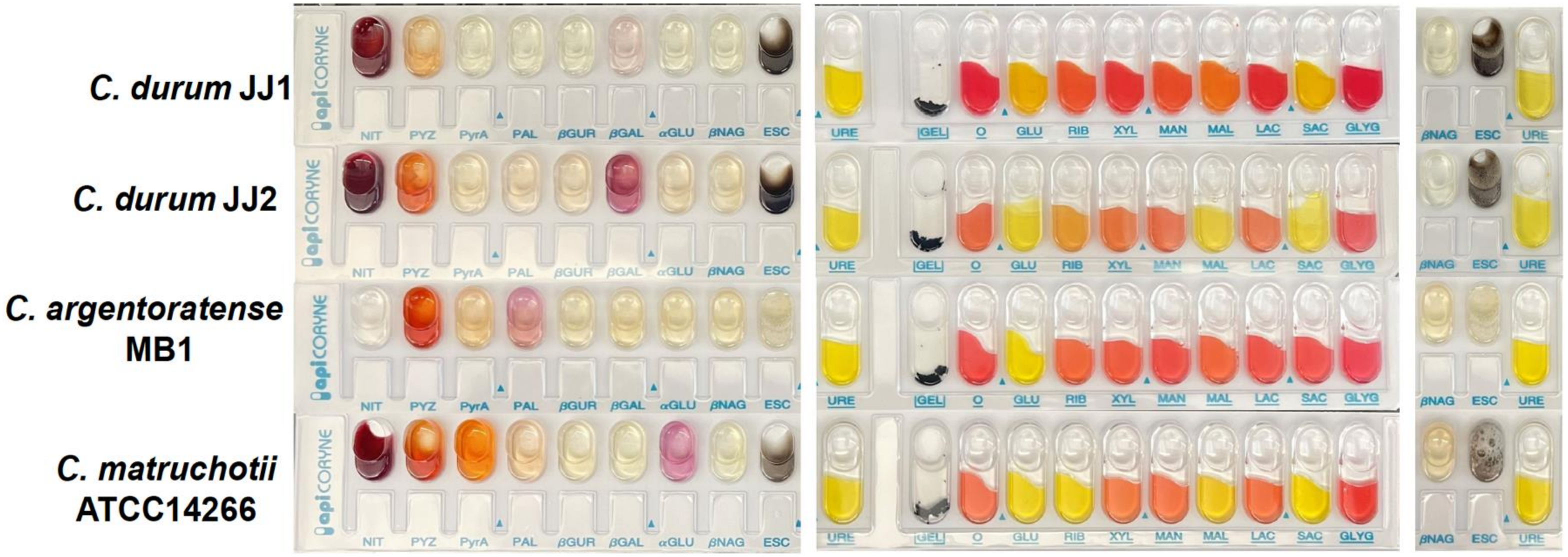


| Strains | NIT | PYZ | PyrA | PAL | βGUR | βGAL | αGLU | βNAG | ESC | URE | GEL | O | GLU | RIB | XYL | MAN | MAL | LAC | SAC | GLYC | CAT |
|---|---|---|---|---|---|---|---|---|---|---|---|---|---|---|---|---|---|---|---|---|---|
| *C. durum* JJ1 | 4+ | 1+ | – | – | – | – | – | – | 3+ | – | – | – | 4+ | 1+ | – | 1+ | 2+ | – | 4+ | – | + |
| *C. durum* JJ2 | 4+ | 3+ | – | – | – | 2+ | – | – | 4+ | – | – | – | 4+ | 2+ | – | – | 3+ | – | 4+ | – | + |
| *C. argentoratense* MB1 | – | 4+ | – | – | – | – | – | – | – | – | – | – | 4+ | – | – | – | – | – | – | – | + |
| *C. matruchotii* ATCC14266 | 4+ | 3+ | 4+ | – | – | – | 2+ | – | 2+ | – | – | – | 4+ | 4+ | – | – | 3+ | – | 4+ | – | + |

Supplemental Figure 2: Biochemical analysis of 4 *Corynebacterium* isolates using Biomeriux API® strips. A. Biochemical tests for the following enzymatic reactions: nitrate reduction (NIT), pyrazinamidase (PYZ), pyrrolidonyl arylamidase (PyrA), alkaline phosphatase (PAL), β-Glucuronidase (β-GUR), β-Galactosidase (β-GAL), α-Glucosidase (α-GLU), N-Acetyl-β-glucosaminidase (βNAG), β-Glucosidase (ESC). Color change indicates positive reaction. B. Biochemical tests for enzymatic reaction of urease (URE), hydrolysis of gelatin (GEL), and fermentation of sugars: glucose (GLU), ribose (RIB), xylose (XYL), mannitol (MAN), maltose (MAL), lactose (LAC), sacrose (SAC), and glycogen. Yellow/orange color indicates positive reaction, red color indicates negative reaction, O cupule used as negative control. C. Hydrogen peroxide test on β-Glucosidase (ESC). Formation of bubbles indicates positive reaction.
